# Integration of publicly available datasets reveals pesticide-associated genetic variation in a wild non-target animal

**DOI:** 10.1101/2025.08.20.671246

**Authors:** Tiffany Scholier, Daniel Guignard, Marissa B. Kosnik

**Affiliations:** Department of Environmental Toxicology, Swiss Federal Institute of Aquatic Science and Technology, Eawag, 8600 Dübendorf, Switzerland

**Keywords:** pesticides, adaptation, resistance, genetic diversity, nervous system function, wild boar

## Abstract

Pesticide use is ubiquitous. While pesticide resistance in target species such as insects is a well-known phenomenon, it is less known how pesticide exposures affect the genetics of non-target species. Here, we demonstrate the research value of integrating existing datasets to study the relationship between chemicals and biodiversity from the genetics perspective by analyzing spatial pesticide concentration data and genetic data from Europe that were not initially generated for toxicological purposes. Specifically, we found associations between environmental pesticide concentrations and genetic variation (*i.e.,* single nucleotide polymorphisms, SNPs) in a non-target species, the wild boar. Gene Ontology (GO) terms and biological signaling pathways related to nervous system function were overrepresented for the impacted genes, especially for insecticides and fungicides. Further, we found that external studies characterizing the molecular mechanisms of these pesticides revealed the same pesticide – GO relationships as those observed in our analysis. Our results show that pesticide-induced genetic changes may not be constrained to target species but may also have an impact on the long-term viability of other wildlife populations. We therefore encourage further research into this area and collaboration across disciplines to develop more comprehensive designated datasets for such analyses. This finding also urges the consideration and inclusion of chemical exposure into conservation protection strategies and greater consideration of genetic impacts from chemicals in environmental risk assessment.

## 1. Introduction

Chemical pollution is one of the three largest drivers of biodiversity loss (Jaureguiberry *et al*., 2022; Keck *et al*., 2025), ahead of climate change and invasive species. While these two latter processes are widely recognized as important stressors within the fields of ecology and conservation genetics, pollution is often overlooked (Sigmund *et al*., 2023). This knowledge gap is concerning as pesticides have been shown to negatively impact important fitness traits (*e.g.,* growth, reproduction) in non-target organisms (Wan *et al*., 2025). Through interference with these physiological processes, pesticides can also be expected to steer natural selection and impact the genetic diversity and genetic composition of wild non-target populations.

Direct toxicity effects and strong selection forces induced by pesticide exposure are expected to decrease the genetic diversity of individuals and populations by removing individuals from the gene pool (through higher mortality and reduced fecundity and mobility) and by eliminating susceptible alleles. Some studies confirm this proposed pattern, as for example, (Siddique, Shahid and Liess, 2024) detected a decrease in genetic diversity between and within populations of a macroinvertebrate species for streams with high pesticide concentrations in comparison to streams with low pesticide concentrations. In contrast, neutral (Gouin *et al*., 2019) and positive (Whitehead *et al*., 2003) associations have been described as well. However, these results have been largely explained by co-occurring biogeographical and evolutionary processes (Whitehead *et al*., 2003), which demonstrates that genetic effects of pesticide exposure in the wild do not happen in isolation.

The existence of pesticide adaptation is a well-known phenomenon, as combatting resistance in target pest species is one of the main concerns within the agricultural sector (Mallet, 1989). However, pesticides are freely released into the environment inevitably exposing more species than the target species (Wan *et al*., 2025), making it very likely that genetic changes following pesticide exposure is a common feature among wildlife. Surprisingly however, research on the genetic effects of environmental pollutants on wild animals besides the target species are relatively rare, especially in the terrestrial environment. One probable reason for the low number of existing studies performed on (terrestrial) wild species is the large requirement in terms of time and funding investment. As an alternative to de novo sampling, computational approaches that can capitalize on the large volumes of already published data have been proposed (Kosnik, Schuwirth and Rico, 2024).

Our aims for this study were two-fold. First, we wanted to test whether we could uncover any associations between pesticide concentrations and genetic diversity (estimated by individual-based and population-based heterozygosity values) and genetic composition (through changes in the allele frequencies of multiple single nucleotide polymorphisms (SNPs)) in a wild non-target species to assess the potential loss of susceptible alleles and to detect a potential adaptive evolutionary response to pesticides. Secondly, we wanted to showcase the research value of repurposing existing datasets and promote computational methods to increase the speed and potential for generating new knowledge of chemical impacts on biodiversity in the context of evolutionary toxicology (Bickham, 2011) and beyond. Specifically, we applied a data-driven method based upon the concept of landscape genetics where we combined genetic variant data (SNP data) for a large terrestrial mammal (wild boar; *Sus scrofa*) that was generated for population genetics studies (Iacolina *et al*., 2016, 2018; de Jong *et al*., 2023) with a disconnected dataset of spatialized pesticide concentrations in the environment (Pistocchi *et al*., 2023). This integrated dataset enabled testing for associations between environmental pesticide concentrations (as a proxy for exposure) and impacted gene functions in a wild non-target organism.

## 2. Material & Methods

### 2.1. Data acquisition

The wild boar SNP data (individuals *n* = 306; SNP *n* = 47,158; (Iacolina *et al*., 2016, 2018; de Jong *et al*., 2023)) and the concentration data (water) for 148 pesticides (Pistocchi *et al*., 2023) were sourced from already published studies. The wild boar data consisted of sampled individuals (2003 – 2010 (∼30% of individuals) and from 2008 – 2018 (∼70% of individuals)) from multiple countries across the European continent for the purpose of studying population structure of wild boar and introgression with the domesticated pig (coordinates of sampled individuals were included). The environmental pesticide data was modelled for EU countries (2015) from *e.g.,* application data by pesticide per country and land use (Pistocchi *et al*., 2023). Therefore, in the absence of any other spatialized data describing pesticide concentrations before 2015, we assume that the pesticide concentrations for 2015 represent average exposure rates that would have been similar enough over time to influence the genetics of the wild boar in a directional way. This is supported by the stable pesticide use in Europe over the past three decades (FigS1, Hader *et al*., 2022; FAO, 2026). To help ensure the biological relevance of this assumption, we only used 121 out of 148 pesticides based on the year of initial application to increase the likelihood of detecting potential evolutionary change due to multigenerational exposure (Year of introduction set before 2000 based on the Pesticide Properties Database (https://sitem.herts.ac.uk/aeru/ppdb/)). Finally, while wild boar are terrestrial mammals, they congregate around water bodies and interact heavily with water through drinking and wallowing (Eckert, Keiter and Beasley, 2019; Brooks et al., 2020) and have even been found to affect water quality themselves due to such high levels of interaction (Hampton et al., 2006; Bolds et al., 2021). Therefore, we consider oral pesticide exposure through water to be a reasonable route for this species. Wild boar coordinates were available as spatial points covering 18 countries and the pesticide data were available as polygons covering 16 of the same countries. For technical details about *e.g.,* the wild boar sample collection or the pesticide concentration modelling, readers are referred to the corresponding publications for the original data sources.

For each pesticide, its type (herbicide, fungicide, insecticide, or other (e.g., molluscicide)) was recorded by using the following online resources: HRAC (Herbicide Resistance Action Committee, https://hracglobal.com/tools/classification-lookup), FRAC (Fungicide Resistance Action Committee, https://www.frac.info/fungicide-resistance-management/by-frac-mode-of-action-group) and the IRAC (Insecticide Resistance Action Committee, https://irac-online.org/mode-of-action/classification-online/) supplemented with the Alan Woods Compendium of Pesticide Common Names (http://www.bcpcpesticidecompendium.org/introduction.html).

Data for conditional variables were retrieved from Köppen-Geiger maps v.3 for climate zones 1991-2020 (111km x 111km, (Beck *et al*., 2018)), JRC-GEOSTAT 2018 maps (1 x 1 km spatial resolution, (Batista e Silva, Dijkstra and Poelman, 2021)) for number of residents/human population density and CORINE Land Cover 2018 maps (100 x 100 m spatial resolution, (Copernicus Land Monitoring Service, 2018)) for agricultural percentage. The largest available resolution for climate data was chosen to ensure that most sampling locations had either one assigned climate zone or at least a clear major assigned climate zone in a 10 km radius.

### 2.2. Merging of datasets

Within PLINK v.1.90b7.2 (Chang *et al*., 2015; Purcell and Chang, 2023), the genetic dataset was filtered to only include SNPs that were at least 90% genotyped and had a minor allele frequency of at least 1%, converted to the raw format with the *recode* command and imported into R v.4.3.2 (R Core Team, 2023). To reduce the introduction of bias, the missing genotype data were substituted with the most common allele type present in the population where the individual boar was sampled (either homozygote for the major allele, homozygote for the minor allele or heterozygote) resulting in the *initial genotype matrix* (individuals *n* = 306; SNP *n* = 40,700). Whenever the number of allele types was equal, missing genotypes were coded as heterozygotes and when all genotypes for a given SNP were missing for a given population, the population was assigned the most common allele type present across all populations. Imputation of missing genotype data is a frequently used method in genetics studies (*e.g.,* (Forester *et al*., 2018; Klein *et al*., 2024)).The mean modeled environmental pesticide concentration within the assumed home range of wild boar was estimated by calculating the mean concentration value of each pesticide present in their home range. Specifically, each pesticide-specific raster was polygonized and a buffer zone of 4 km radius was created around each of the individual coordinates to approximate the home range of wild boar (∼50 km^2^ as an average of home range sizes mentioned in (Miettinen *et al*., 2023) and references therein) resulting in multiple segments with different concentration values within the 4km buffer zone (an illustrative example can be found in the Supplementary Information as Figure S2). The average concentration within the buffer zone range for each of the 121 pesticides was calculated resulting in the estimated pesticide exposures per individual wild boar. Similarly, human population density and agricultural percentage were calculated within the radius for each individual wild boar (the area size of land use categories assigned as “agricultural areas” by CORINE was divided by the total area size within the radius). Additionally, climate categories were assigned to each individual based on a 10 km radius.

To ensure that the selected 4 km radius was not heavily influencing our results, the same analysis was repeated for pesticide concentrations, agricultural percentage and human population density for a smaller (3 km radius) and (5 km radius) larger home range for each wild boar individual (climate zones were always calculated for a 10km radius). This resulted in three different versions of the *final pesticide metadata file* based on the chosen radius.

### 2.3. Patterns in genetic diversity

The *initial genotype matrix* was subset for individuals for which pesticide data was available (pesticide data for Bosnia and Herzegovina and Serbia were missing) resulting in the *final genotype matrix* for further analysis (individuals *n* = 282; SNP *n* = 40,700).

Individual-level heterozygosity values (H_IND_) were calculated from the *final genotype matrix* by dividing the number of heterozygous loci per individual by the total number of loci. To make an estimation of the population-level genetic diversity metrics (observed heterozygosity (H_O_), expected heterozygosity (H_E_)), the *final genotype matrix* was filtered to only include individuals that share their geographical coordinates with at least three other individuals resulting in the *final population genotype matrix* (individuals *n* = 199; SNP *n* = 40,700; populations *n* = 30 with minimum 4 individuals per population). The standard formulas for both metrics were applied per population and averaged across all 40,700 SNPs:

- **Averaged Observed Heterozygosity** (percentage of individuals in a population that are heterozygous for a given SNP, averaged across all SNPs)

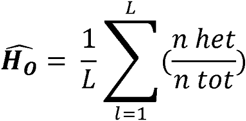

With:

- *L* = total number of SNPs
- *n het* = number of heterozygous individuals for SNP *l*
- *n tot* = number of total individuals for SNP *l*

- **Averaged Expected Heterozygosity** (percentage that two alleles randomly drawn from the gene pool are different (*i.e.,* heterozygous) assuming Hardy-Weinberg equilibrium for a given SNP, averaged across all SNPs)

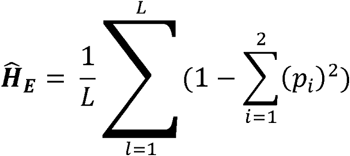

With:

- *L* = total number of SNPs
- *p_i_* = frequency of allele *i* for SNP *l*

For each of the three genetic diversity metrics, two linear models were constructed with the genetic diversity metric as the response variable and latitude, longitude, agricultural percentage, and human population density as explanatory variables (climate zone was excluded for consistency with the RDA models; see Section 3.2). The first model included pesticide concentration while this variable was excluded in the second model. We used ANOVA to test for a difference in explanatory power between the two models which would explain whether pesticide concentration significantly contributed to the variation in heterozygosity levels. Specific adjustments to the linear models were the inclusion of the quasibinomial distribution (for the H_IND_ as this responds to proportional value) and the log-transformation of the genetic metric (for H_O_ and H_E_). The relationship between the concentrations of all significant pesticides and the genetic diversity metric was tested by subtracting the explained variation (R^2^) between the two models. To reduce the chance of outlier populations heavily impacting the results, the linear model was only run for a certain pesticide if it had data available for at minimum 5 populations.

### 2.4. Identifying unique SNP-pesticide links

Significantly correlating SNPs were identified per pesticide and mapped onto the *Sus scrofa* reference genome to match intragenic SNPs with their respective genes. Specifically, 121 redundancy analyses (RDA) were applied with the *final genotype matrix* as the response variable, one out of the 121 pesticide concentrations as the explanatory variable and latitude, longitude, agricultural percentage, human population density, and climate zone as conditional variables. Multiple test corrections were controlled by using the false discovery rate (FDR) (*adjusted p-value* < 0.05). Significant SNPs were remapped onto the latest version of the *Sus scrofa* reference genome (Sscrofa11.1), with the use of the SNP identifiers and the Illumina PorcineSNP60 v2.0 data file (downloaded from https://emea.support.illumina.com/array/array_kits/porcinesnp60_dna_analysis_kit/downloads.html) since the original map and ped files were based on an earlier version of the *Sus scrofa* reference genome (Sscrofa10.2). Next, a gene annotation file was downloaded from Ensembl ( https://ftp.ensembl.org/pub/release-113/gtf/sus_scrofa/) to match each intragenic SNP with its corresponding protein-coding gene. The resulting dataset included links between SNPs, pesticides and genes and is referred to as the *initial SNP-pesticide dataset*.

To assess multicollinearity among explanatory variables and the overall significance and explanatory power of each RDA model, we calculated variance inflation factors (VIFs), performed permutation tests, and calculated adjusted R² values. When a variable exceeded the predefined VIF threshold, it was removed and the model was refitted before subsequent analyses.

### 2.5. Sensitivity analysis

To apply a robustness check to exclude associations that are potentially driven by factors other than pesticide concentrations, a sensitivity analysis was conducted where the concentrations of the pesticides were randomly shuffled between individuals before the initial RDA analysis was repeated (iterations per pesticide *n* = 100). Each SNP-pesticide link that occurred multiple times in the shuffled SNP-pesticide datasets was removed from the *initial SNP-pesticide dataset*. This resulted in the *final SNP-pesticide dataset*.

### 2.6. CTD data curation

The Comparative Toxicogenomics Database (CTD; (Davis *et al*., 2025)) is a public database that curates chemical, gene, pathway, and disease-related relationship data from the literature. All CTD data were downloaded from the 30/06/2025 release of CTD. Data on chemical – gene interactions reported in studies for *Sus scrofa* were collected and, because data on chemical – gene interactions for *Sus scrofa* are limited in the literature, associations reported for all other species were also collected to provide additional comparison between reported pesticide – gene interactions. To match Gene Ontology (GO) terms and pathways in our analysis to specific toxicity endpoints (described below), classes for pathways and GO terms were set based on the “Disease-pathway” associations and “Phenotype (GO)-Disease Inference Network” files from CTD. To ensure that the assignment of GO and Kyoto Encyclopedia of Genes and Genomes (KEGG) pathway terms to disease classes was not arbitrary, we applied a systematic, rule-based approach using CTD’s “MEDIC disease vocabulary” classification system which uses the National Library of Medicine’s Medical Subject Headings to classify human diseases into distinct categories related to disease type (*e.g.,* nervous system disease, cancer, cardiovascular disease). Briefly, each file contains a set of GO terms/pathways associated with different human diseases. For each term, we identified the corresponding disease category (based on CTD’s Medic disease vocabulary), and then determined how many of the associated diseases per term fell into each MEDIC category. From this, we identified the most common disease category per GO term/pathway. For GO terms that serve as nodes in the ontology, the primary disease category for all the diseases within that node was used. In cases where a GO term corresponded to multiple disease categories or was unclassified, the class was set as “Other” (*e.g.,* “transferase complex”).

### 2.7. Characterizing potential toxicity mechanisms

All associations between Reactome pathways, Gene Ontology terms and wild boar genes (*Sus scrofa*) and human genes (*Homo sapiens*) were retrieved from the DAVID knowledgebase (Huang, Brad T Sherman and Lempicki, 2009; Huang, Brad T. Sherman and Lempicki, 2009) on 22/07/2025. KEGG pathways and associated gene sets for both *Sus scrofa* and *Homo sapiens* were collected on 10/07/25 using the get_gene_sets_list function from pathfindR package (Ulgen, Ozisik and Sezerman, 2019). GO and pathway enrichment was done for our landscape genetics associations (based on all pesticide – gene associations, or separately per individual pesticide and per pesticide class) and CTD pesticide – gene associations for humans. For gene set enrichment, overrepresentation analysis was done using the hypergeometric test with a requirement of a minimum of three genes overlapping between comparison datasets (*adjusted p* < 0.05; multiple-test corrections using FDR). The background was set as all unique genes represented across the GO, KEGG and Reactome annotation datasets per species tested, and only GO terms/pathways with at least five genes and at most 1000 genes associated were included. The lower threshold was set to ensure enough genes were present for each GO term/pathway to avoid significant results by chance or associations to poorly characterized mechanisms. The upper threshold was set to exclude generic pathways/terms described by huge numbers of genes (*e.g.,* “Signal transduction”). Enrichment analysis was done for all pesticides at once, per pesticide, and per pesticide class. For enriched GO terms/pathways, the corresponding MEDIC disease category was used to classify the term as more associated with *e.g.,* nervous system disease endpoints or cancer.

### 2.8. Data Analysis and Visualization

Analyses were done using R v.4.6.1 (R Core Team, 2023). Packages include adegenet v.2.1.10 (Jombart, 2008), rsq v.2.7 (Zhang, 2024), vegan v.2.7.5. (Oksanen *et al*., 2026), qvalue v.2.44.0 (Storey *et al*., 2026), rtracklayer v.1.72.0 (Lawrence, Gentleman and Carey, 2009), data.table v.1.18.4 (Barrett *et al*., 2026) and openxlsx v.4.2.9. (Schauberger and Walker, 2026). Figures were made with ggplot2 v.4.0.3 (Wickham, 2016), cowplot v.1.2.0 (Wilke, 2025), ggtext v.0.2.0 (Wilke and Wiernik, 2026), ggrepel v.0.9.8 (Slowikowski, 2026), patchwork v.1.3.2 (Lin Pedersen, 2025) and igraph v.2.3.3 (Csárdi *et al*., 2026) R packages. Spatial analyses were done using sf v.1.1.2 (Pebesma and Bivand, 2023), terra v.1.9.34 (Hijman, Brown and Barbosa, 2026) and exactextractr v.0.10.1 (Baston, 2025) R packages..

## 3. Results

To the best of our knowledge, there are no public resources that contain both chemical exposure measurements and population genetics data. To combat this challenge, we collected and integrated existing, disparate datasets for genetic variants (total SNP *n* = 40,700, total individuals *n* = 282) for a large terrestrial mammal (wild boar; *Sus scrofa*,) and geospatial data for environmental freshwater pesticide concentrations (as a proxy for exposure) from different sources (Iacolina *et al*., 2016, 2018; de Jong *et al*., 2023; Pistocchi *et al*., 2023). Our integrated starting dataset comprised 40,700 SNPs from 282 individuals and environmental concentration data for 121 pesticides across Europe together with data on potential confounding environmental variables (*e.g.,* agricultural percentage of land area) (Figure 1).

**Figure 1.**
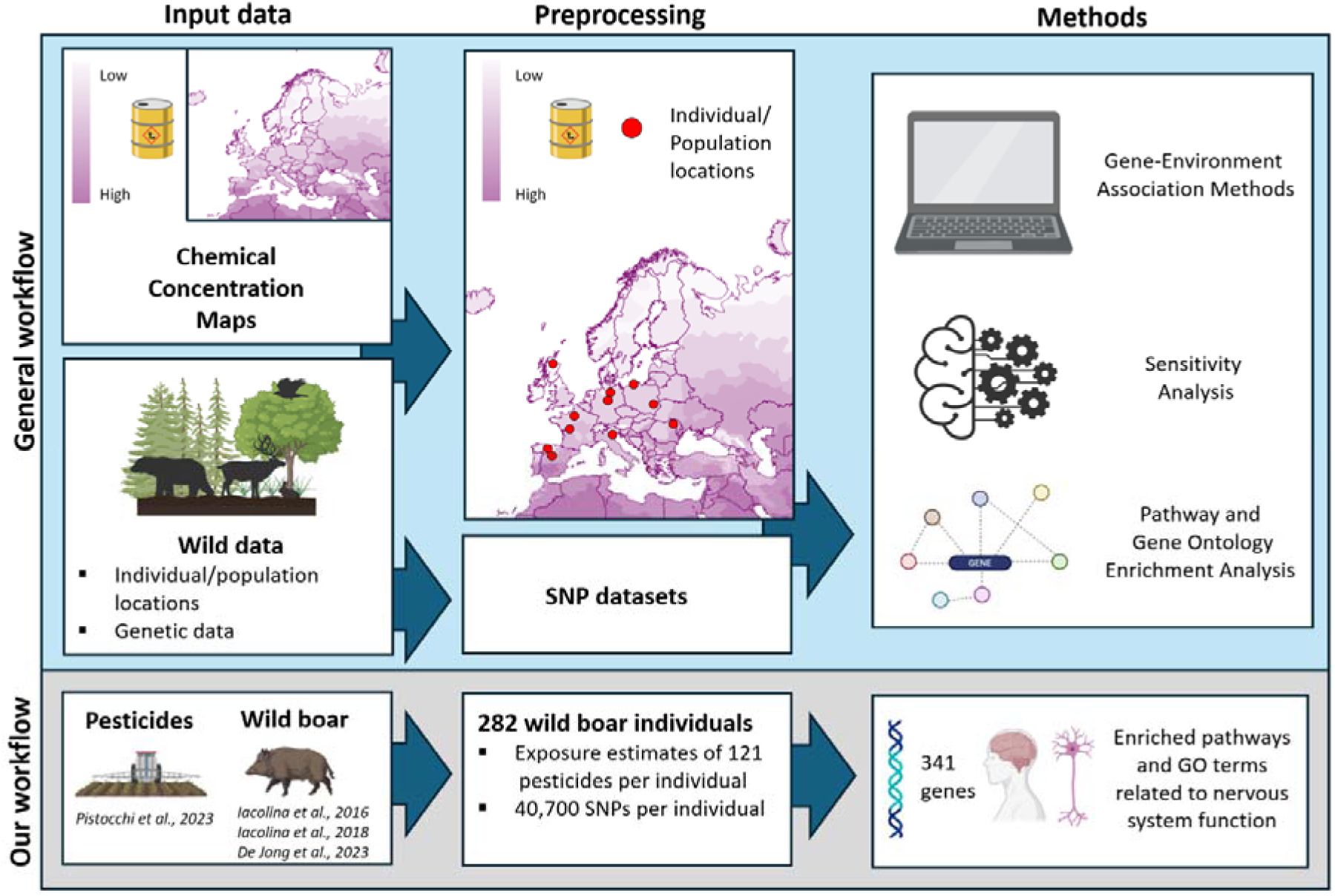
Landscape genetics workflow. The figure summarizes the data inputs and the processing steps taken in this study that we propose as a workflow to computationally characterize chemical impacts on the genetics of wild populations, based on the concept of landscape genetics. In short, wild genetic data and chemical concentration maps from independent sources are integrated using individual sampling coordinates to identify associations between chemical exposure and genetic variation. From there, genes with genetic variability can be assessed for function using gene enrichment to evaluate the possible mechanisms of chemical toxicity and/or adaptation. Our study followed this approach using pesticides and wild boar. Illustrations were created with BioRender.com.

### 3.1. Associations between pesticide concentrations and genetic diversity are pesticide dependent

To assess whether pesticide concentrations associate with wild boar genetic diversity, we first estimated individual-level heterozygosity values (H_IND_) (individuals *n* = 282; SNP *n* = 40,700) and two population-levels metrics (observed heterozygosity (H_O_) and expected heterozygosity (H_E_)) across 30 populations (individuals *n* = 199; SNP *n* = 40,700; populations *n* = 30 with minimum 4 individuals per population).

Individual heterozygosity levels varied between 10% (West-Sardinia) and 45% (Estonia) with an average of 23% across all individuals. Out of 121 pesticides, 40 pesticide concentrations were significantly associated with variation in individual-level heterozygosity with the majority (29 out of 40) demonstrating negative correlations. The explanatory variation explained per chemical ranged between 0.9% (amidosulfuron) and 21.2% (diclofop) with an average of 3.5% across all chemicals (Table S1).

In terms of population-level genetic diversity, the values ranged from 14.1 (H_E_) – 14.7% (H_O_) for West-Sardinia) to 34.8%-35.1 % (both H_E_ and H_O_ for Estonia) with an average of 21.1 (H_E_) – 22.9% (H_O_) across all populations. While 116 out of 121 chemicals had enough data points to continue the analysis (5 chemicals did not have data for the minimum set value of 5 populations), only 7 and 5 chemicals significantly explained variation in H_E_ and H_O_, respectively (Table S2). Chemicals showed both negative and positive correlations, with MCPB explaining the largest amount of variation for both metrics (H_E_: 68.5%, H_O_: 47.2%).

### 3.2. Different SNP alleles are associated with high and low pesticide concentrations

To detect SNPs that associate with environmental concentrations for each of the 121 pesticides, a redundancy analysis was carried out for each pesticide. Due to high VIF values, climate zone was removed as a conditional variable (Table S3). With the final model (including latitude, longitude, agricultural percentage, and human population density as conditional variables), we found 1,403 significant SNP-pesticide links divided between 50 pesticides (analysis of 71 pesticides did not reveal any significant SNPs). The final RDA models generally explained a small proportion of the genetic variation, with adjusted R² values ranging from 0.13% to 1.77%. Overall, 109 of 121 models were significant based on permutation tests (*p* < 0.05). After removal of climate zone, four pesticide models (Bifenthrin, Carbetamide, Diclofop and Forchlorfenuron) retained VIF values above the predefined threshold and were excluded from subsequent analyses. VIF values, permutation-test results, and adjusted R² values for all models are provided in Table S4.

To remove SNP–pesticide associations that repeatedly occurred under randomized pesticide concentration assignments, we shuffled pesticide concentrations between individuals and identified significant SNPs according to the same workflow mentioned above (iterations *n* = 100). Any overlap between the initial 1,403 SNP-pesticide links and the randomized SNP-pesticide links that occurred multiple times in the sensitivity analysis dataset were removed from the initial dataset as these represented SNPs that may be associated with factors other than the pesticide concentrations. This resulted in the *final SNP-pesticide dataset* comprising 1,238 SNP-pesticides links (514 intra- and 724 intergenic links) for 50 pesticides (see Table S5). All intragenic SNPs were linked to 39 pesticides and 278 unique genes (range of intragenic SNPs found per pesticide: 1 – 75). The list of the pesticide – SNP linkages removed through the sensitivity analysis can be retrieved from Table S6. Only the results obtained using the 4-km radius are discussed in the main text. Corresponding results for the 3- and 5-km radii are provided in Tables S7–S10, and the overlap in associated SNPs among the three radii is shown in Figure S3.

By examining the allele frequency changes in the significant SNPs associated with the five most frequently occurring genes, we found that the significant associations with pesticide concentrations were mostly driven by specific geographic locations rather than exhibiting a widespread pattern across multiple regions (Table S11).

### 3.3. Intragenic SNPs associated with pesticide concentrations are within genes associated with different pesticide classes

To determine the potential biological implications of the detected pesticide-associated genetic variation in wild boar, gene set enrichment analysis (*FDR-adjusted p < 0.05*) was conducted for the 278 genes derived from the landscape genetics analysis with *Sus scrofa* Gene Ontology (GO) terms and KEGG and Reactome pathways.

By assessing the enrichment across all 278 genes independent of their associated pesticide, we identified significant enrichment of 89 Gene Ontology/KEGG/Reactome terms: 25 biological processes (BP, Figure 2A), 33 cellular components (CC, Figure 2B), 19 molecular functions (MF, Figure 2C), 8 KEGG pathways, and 4 Reactome pathways (Figure 2D). More than half of the significantly enriched biological processes (23 out of 25 = 92%), cellular components (25 out of 33 = 76%), molecular functions (15 out of 19 = 79%) and KEGG/Reactome pathways (8 out of 12 = 67%) have terms that could be either linked to nervous system function (mentioning terms “nervous”, “neuro(n)”, “axon”, “synaptic/synapse”, “dendrite” or “potentiation”) or membrane transport-related processes (mentioning terms “transmembrane”, “channel”, “cation/ion” and “transport”. The three terms with the largest gene overlap (with the FDR-adjusted *p-value* used to distinguish terms with equal overlap) were cell motility, monoatomic ion transport and cell migration for BP, neuron projection, plasma membrane region, and plasma membrane protein complex for CC, monoatomic ion transmembrane transporter activity, inorganic molecular entity transmembrane transporter activity, and channel activity for MF, and human papillomavirus infection, neuronal system, and axon guidance for KEGG/Reactome pathways. The full list of enriched GO/KEGG/Reactome terms and associated genes can be retrieved from Table S12. The enrichment results for the overlapping SNPs between the three radii can be found from Table S13 and Figure S3.

**Figure 2.**
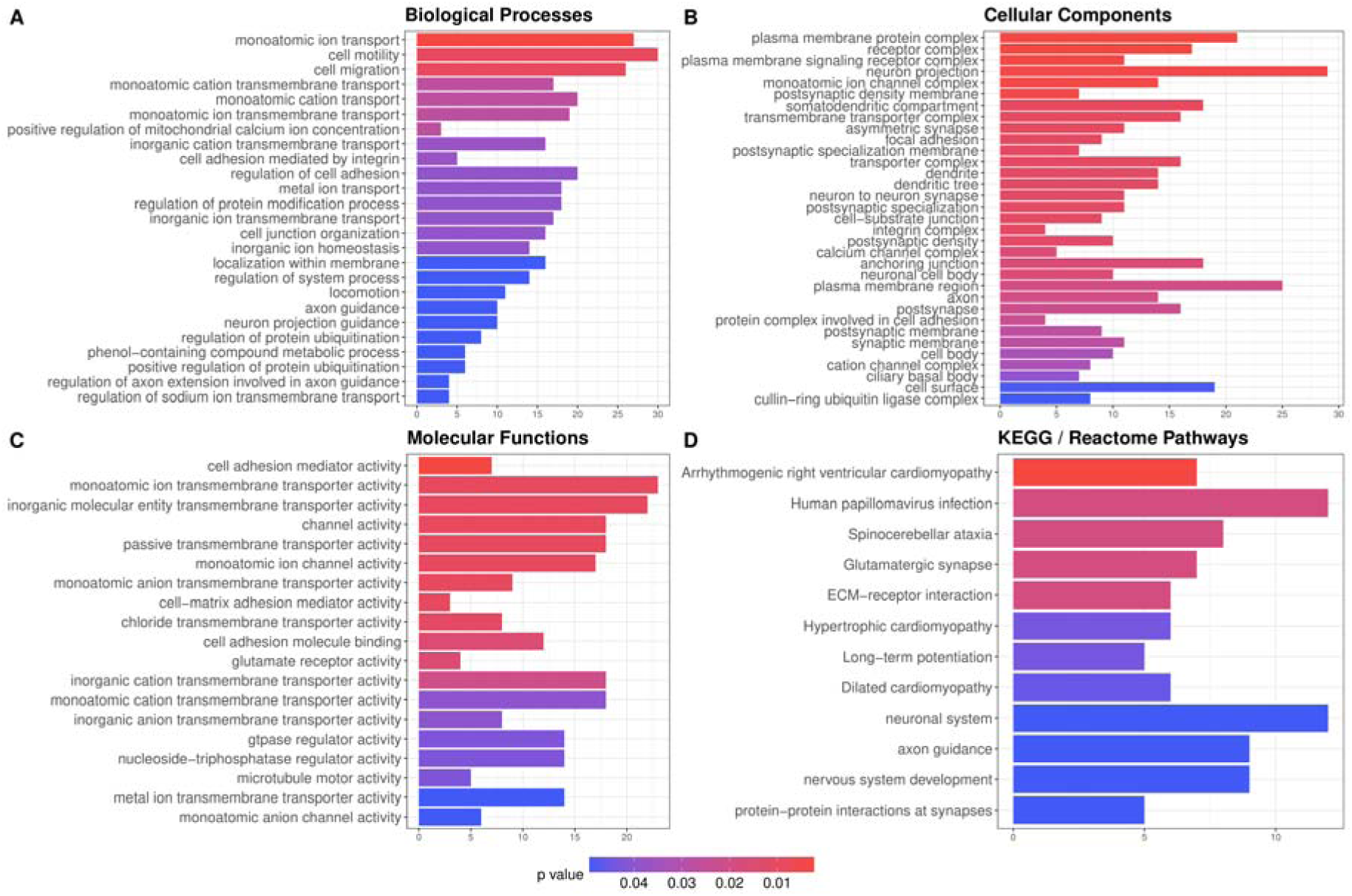
Enrichment of Gene Ontology and pathway terms for *Sus scrofa* genes with SNPs associated with environmental pesticide concentrations. Enriched Gene Ontology terms ((A) biological processes, (B) cellular components and (C) molecular functions) and pathway (KEGG) terms (D) for the 278 wild boar genes with SNPs significantly associated with environmental pesticide concentrations. Results are shown for all genes together, regardless of their associated pesticide. Color = FDR-corrected p-value.

To assess the relative influence of each pesticide in the overall enrichment pattern observed across all 278 genes in the landscape genetics dataset, we conducted gene set enrichment analysis separately for each pesticide using only the genes associated with significant SNPs for that specific pesticide (1 – 70 genes per pesticide). Through this, 14 pesticides had at least one GO or pathway term (KEGG or Reactome) associated, with pyrimethanil, formetanate, and dimethomorph having the most enriched terms (58, 52, and 27 terms, respectively). The full list of pesticide – GO/pathway terms is in Table S14. To assess whether different pesticide classes (with different modes of action) were linked to specific SNPs in genes, we grouped the pesticides according to class before comparing the enriched pathways and GO terms corresponding to nervous system disease for each pesticide type (insecticides, fungicides, herbicides, and other pesticides). Nervous system-related diseases were the most represented disease class in the dataset and were more often linked to insecticides (3 insecticides with 29 enriched terms) and fungicides (6 fungicides with 53 enriched terms) than to herbicides (1 herbicide with 1 enriched term) and other pesticides (2 pesticides with 4 enriched terms).

Additionally, we conducted gene set enrichment analysis separately for each of the four pesticide classes using only the genes associated with that class (32–160 genes per pesticide class). Insecticides and fungicides showed a higher proportion of enriched terms related to nervous system disease than herbicides and other pesticides: 49% for insecticides (21 of 43 terms), 48% for fungicides (28 of 58), 20% for herbicides (1 of 5), and 25% for other pesticides (2 of 8; Table S15, Figure S4). These class-level differences were accompanied by substantial class specificity in the underlying genetic associations, with 50% of fungicide-associated SNPs, 57% of herbicide-associated SNPs, and 61% of insecticide-associated SNPs being unique to that pesticide class (in contrast to only 13% of the pesticides grouped in the category “Other”).

### 3.4. External datasets support that pesticides implicate some of the same genes, pathways, and GO terms in mammals

To determine if the GO terms and pathways enriched for different pesticides based on the SNPs/genes identified using the landscape genetics approach have external evidence supporting our predicted associations, we looked at existing external data on pesticide – gene interactions for *Sus scrofa* in the Comparative Toxicogenomics Database (CTD, primarily *in vitro* studies). Two pesticides from our landscape genetics study were reported to have known chemical-gene interactions in *Sus scrofa*: chlorpyrifos-methyl and malathion. While none of the genes overlapped between the Sus scrofa CTD data and our landscape genetics results for these specific two pesticides, this comparison was based on relatively small gene sets: the two pesticides implicated 14 unique genes in our landscape genetics analysis and 19 unique genes in CTD (of which only 8 were in the initial input SNP dataset).

Because CTD has comparatively less chemical – gene interactions data for *Sus scrofa* than for humans, we also compared enriched GO and pathway terms between pesticide – gene interactions curated in CTD and those identified in our landscape genetics study that are not specific to *Sus scrofa*. For humans, 30 pesticides from our landscape genetics dataset had chemical – gene interactions curated in CTD, which provides external support that these pesticides have been found to affect gene activity in mammals. Further, 15 pesticide – gene associations that were found in our landscape genetics approach overlap with the pesticide – gene interactions reported for humans and other commonly-studied species in CTD, which provides explicit support for the pesticide – gene associations captured in our landscape genetics analysis (Table S16). Finally, 16 GO and pathway terms were implicated by the same 6 pesticides in both CTD and our landscape genetics dataset (Cypermethrin, Dimethomorph, Folpet, Malathion, Pyrimethanil and Thiram) with malathion and thiram implicating four exact pesticide–gene–GO/pathway overlaps (ITGB8, DDB1, GHR and ITGA1). Within these 16 GO and pathway terms, both nervous system-related (e.g., regulation of membrane potential) and cell regulation-related (e.g., cellular homeostasis) terms were highlighted (Table S17). The full network of overlapping terms and pesticides for CTD human data and our wild boar landscape genetics data can be viewed in Figure S5 and the SNPs in genes underlying these GO terms/pathways for all 6 pesticides is demonstrated in Figure 3. Of the landscape-genetic genes contributing to the overlapping GO terms and pathways, 8 were also represented in the human CTD enrichment dataset but were associated with different pesticides, whereas 22 were not represented in the corresponding CTD starting data. These genes therefore provide additional candidates for experimental studies investigating mechanisms of potential susceptibility and/or adaptation to chemical pollution.

**Figure 3.**
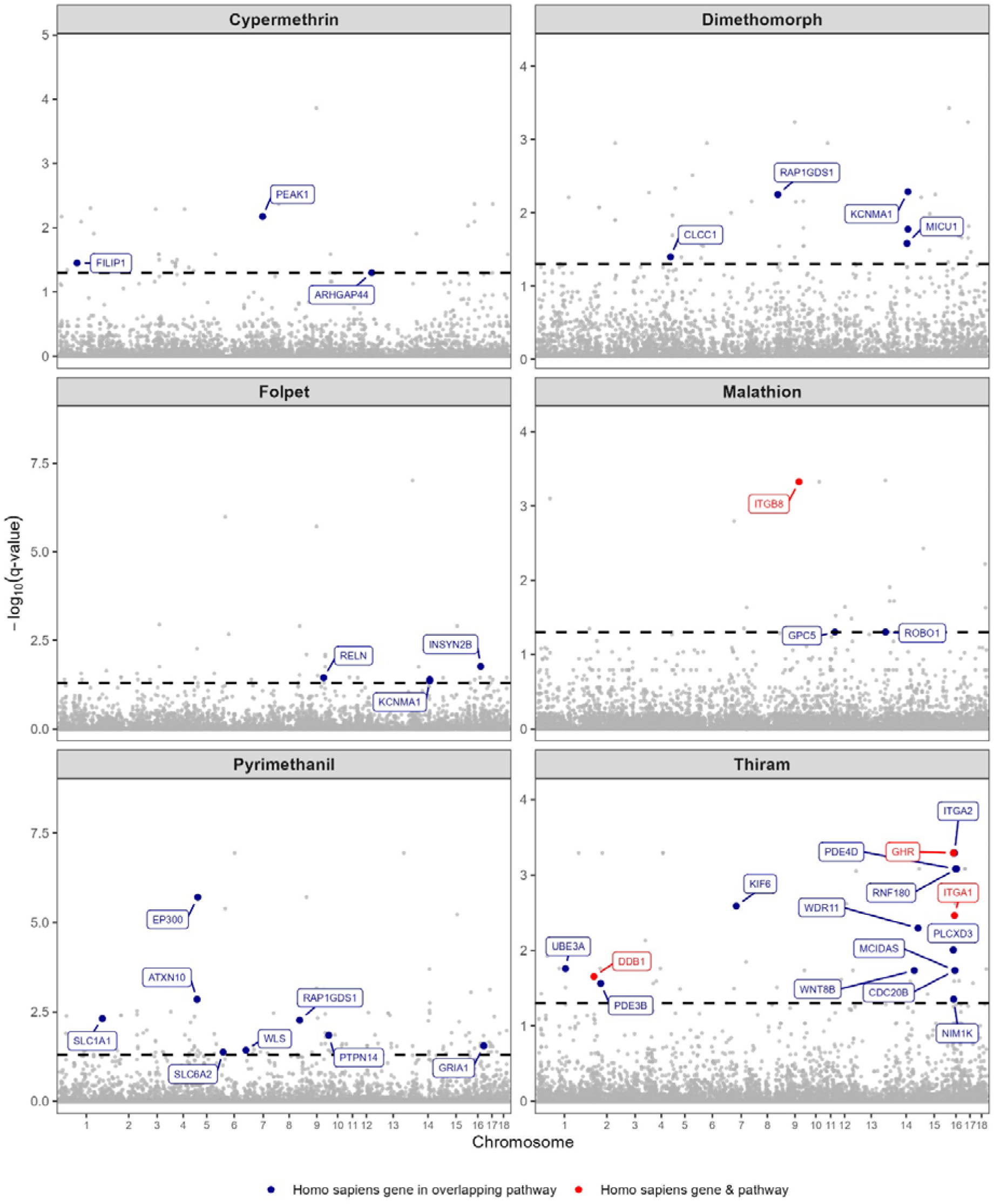
Pesticide-associated SNPs in genes supported by comparison with the Comparative Toxicogenomics Database (CTD). Manhattan plots show SNP associations for six pesticides for which exact genes (red) or enriched GO terms or pathways (blue) identified in the landscape genetics analysis for *Sus scrofa* overlapped with data in CTD for *Homo sapiens*. The dashed horizontal line indicates the significance threshold.

## 4. Discussion

By valorizing existing publicly available datasets, this study demonstrates that higher pesticide concentrations distributed over the European continent coincide with changed allele frequencies of intra- and intergenic SNPs in wild boar (frequency of susceptible alleles decreased in more pesticide-exposed populations). All identified intragenic SNPs were mapped onto 278 genes, which were found to be enriched in terms of nervous system function based on Gene Ontology and pathway analysis. Insecticides and fungicides, as opposed to herbicides and other pesticides, acted as the main driver behind this result. Although these results are based on correlational data, they raise the important question of whether evolutionary responses to pesticide exposure, especially exposure to insecticides and fungicides, occur in non-target wild species.

Associations between genetic diversity and pesticide concentrations varied depending on the pesticide and the genetic diversity metric. At the individual level, significant associations were predominantly negative, with 29 of the 40 associated pesticides showing decreasing heterozygosity with increasing pesticide concentrations. In contrast, relatively few pesticides were associated with population-level genetic diversity, and these associations varied in direction depending on the pesticide. The predominantly negative associations observed at the individual level are consistent with the hypothesis that environmental stressors such as pesticide exposure may be associated with reduced genetic diversity (Bickham *et al*., 2000). For example, sustained exposure could increase mortality and thereby reduce the effective number of individuals contributing to subsequent generations. The absence of a similarly clear pattern at the population level may reflect differences in how quickly individual- and population-level genetic diversity respond to environmental pressures. Changes in population-level genetic diversity can accumulate across generations (Nowak *et al*., 2009), and there may therefore have been insufficient time for pesticide-associated effects to be to become readily detectable at the population level. Methodological factors could also contribute to the differences between metrics, including the considerably smaller sample size at the population level (30 populations compared with 282 individuals) and potential biases associated with the SNP panel, which may not fully capture genome-wide genetic diversity.

The observed associations between hundreds of wild boar SNPs and pesticide concentrations suggest that, if some of these associations reflect genetic responses to pesticide exposure, such responses may involve changes across many genes (i.e., polygenic resistance, as was proposed for glyphosate resistance in a weed plant (Kreiner et al., 2021)). While primary adaptations in a large number of genes may be unlikely as many pesticides are designed to have a very specific mode of action with deliberate target genes, it could be possible that secondary compensatory adaptations evolved to the direct and primary genetic adaptations, as a response to fitness trade-offs and/or to correlating environmental variables (Whitehead et al., 2017). Interestingly, we found genes associated with pesticide concentrations to be more likely linked to nervous system-related GO terms and pathways. Comparison across the four pesticide types (insecticides, herbicides, fungicides, and other pesticides) revealed that nervous system disease-related terms and pathways were particularly enriched for insecticides and fungicides. Further, approximately 50% of the SNPs associated with each pesticide class were unique to that class rather than shared among classes. Thus, differences in functional enrichment among pesticide classes cannot be explained solely by the same SNPs being repeatedly associated with multiple classes. The pattern observed for insecticides is consistent with the known neurotoxicity of many insecticides, which are designed to disrupt nervous system signaling pathways that can be conserved across animal taxa (Ganie et al., 2022) while detecting the same pattern for fungicides suggests that fungicides can also impact non-target organisms (Zubrod *et al*., 2019). Previous studies in humans have similarly linked pesticide exposure to neurological outcomes. For example, other data driven studies have reported associations among regional pesticide use, neurodegenerative disease incidence, and implicated genetic variants (Kosnik, Antczak and Fantke, 2024), while there is growing interest in the potential role of fungicides in neurological disease development (Lini *et al*., 2024). Together, these patterns suggest that differences among pesticide classes may reflect biological differences in their modes of action, although potential effects of variation in exposure, environmental persistence, pesticide representation in the dataset, and statistical power cannot be excluded.

Some external support for the pesticide-SNP associations identified by the landscape genetics method was found by comparing the pesticide–gene associations in our landscape genetics dataset with those curated from the literature (CTD database). Specifically, two pesticides in our analysis have been previously reported to alter the expression of the same genes, albeit in different vertebrate species. Also, a subset of pesticide – GO terms/pathways links identified using the landscape genetics method were also formed from the CTD data. However, many of the pesticide – gene/GO terms/pathways linkages reported in our analysis or reported in CTD were not found in the other dataset. This is not surprising as the data from CTD was collected from studies conducted for different purposes than the datasets in our analysis, and thus focused on different genes than those included in the landscape genetics dataset. For example, many of the pesticide – gene associations for *Sus scrofa* in CTD came from studies assessing the influence of pesticide exposure on pig development, either *in utero* or in pig embryos *in vitro*, *e.g.,* (Salazar *et al*., 2007; Chen *et al*., 2020; Jiang *et al*., 2021; Bai *et al*., 2023). Therefore, given that many of the associations from the landscape genetics analysis were related to nervous system function GO terms/pathways, it is not surprising that there was limited overlap between genes studied in our dataset and CTD. Further, future studies assessing the impacts of pesticides on non-target organisms like wild boar can use the associations we found to guide analyses. For example, studies of malathion on wild populations can search for possible SNPs in the genes implicated in CTD but that were not included in the SNP array used for the landscape genetics part of our study (*e.g.,* ACHE, CYP1B1, and GPX4) while future lab studies can assess activity in the genes we found through the landscape genetics approach that were not in the CTD studies (*e.g.,* ROBO1, GPC5 and SNX7).

A huge benefit of our analysis is that we harness datasets that were published for other research purposes and increase their utility in the context of new research questions. However, while working with pre-existing data enables cost- and time-effective analyses, it simultaneously poses specific challenges such as 1) the reliance on predicted pesticide concentrations as proxies for multigenerational pesticide exposure, 2) the dependence on the quality and quantity of available input data and 3) multicollinearity between pesticides and other environmental variables. By only including pesticides that were in circulation before the year 2000, we increased the likelihood that the actual pesticide exposure experienced by the wild boar populations over time was sufficient to theoretically induce adaptive responses. While it is still possible some specific pesticide use may not have been consistent over time, pesticide use in Europe as a whole has been relatively stable for the past decades (Figure S1, (Hader *et al*., 2022; FAO, 2026)) and agricultural land use has only decreased by 5% across Europe since 2000 (FAO, 2025). To improve estimates of pesticide exposure in future studies, direct measurements from biological tissues could be obtained through dedicated and invasive field sampling or from existing sources such as hunter associations. Nonetheless, such measurements would still represent current exposure levels at the time of collection and would still serve only as proxies for the exposure levels experienced by previous generations. To truly overcome these limitations, long-term research efforts combining environmental monitoring with repeated measurements of pesticide concentrations in tissues across generations in multiple individuals in multiple locations would be necessary. Another aspect to take into consideration is that the genetic data for this study was based upon a subset of SNPs and thus represents only part of the true genetic variation present in the species. Additionally, the pesticide concentration data was extrapolated from regional application rates rather than direct measurements and was limited to certain geographic regions and certain pesticides. Also, the modelled freshwater concentrations represent only one component of potential pesticide exposure in wild boar (e.g., contaminated soil, vegetation, etc) and therefore serve as proxies of pesticide exposure rather than estimates of total individual pesticide uptake. As such, our significant associations between pesticide concentrations and allele frequencies were mostly driven by certain geographic regions, and further analysis is necessary to discern whether this pattern reflects biologically meaningful processes such as localized adaptation or if it is simply a result of input data bias. The use of whole genome sequencing data and the inclusion of more individual data points and broad-coverage pesticide concentration data (from different potential exposure sources) if these datasets become available in the future could help reduce these potential false negative (or false positive) results by making the analysis more robust and revealing geographically widespread adaptation patterns that can be cross-validated. Lastly, an unavoidable challenge in studying the impacts of pesticides is the disentanglement of effects between individual pesticides as many pesticide concentrations naturally correlate with one another (and other environmental variables) which makes it impossible in this study to confidently assign genetic effects to single compounds. This is important as exposure to a single chemical, a combination of chemicals or a combination of different stressors (*e.g.,* parasite and viral load) can all result in different fitness and adaptation outcomes (Hua *et al*., 2017). Potential environmental and spatial confounding was partly controlled for in our study by using a conditional RDA model with the inclusion of agricultural land use, human population density, latitude, and longitude as conditional variables. However, not all potential correlating variables could be tested, nor would pesticides correlating with each other have been disentangled. By expanding this analysis to include more genetic variants, it could be possible that the association between specific SNPs/genes and pesticide modes of action can be better characterized to help disentangle which genetic adaptation can be attributed to specific pesticides (*e.g.,* associations between AChE inhibitors and SNPs/genes in cholinergic signaling pathways could be assumed to be better associated than correlating pesticides that target the cell wall).

To the best of our knowledge, this study is the first to demonstrate an association between environmental pesticide concentrations and genetic variation in a non-target wild species, suggesting potential selective processes related to pesticide exposure. While these potential adaptations may help organisms withstand pesticide toxicity, they may also come at a cost as resistance to (chemical) stressors often carries trade-offs that can impact the ability to withstand other stressors (Pedra et al., 2004; Cutti et al., 2021) and/or long-term fitness (Shirley and Sibly, 1999; Jansen et al., 2011). Maintaining genetic diversity (encompassing both resistant and susceptible alleles) is therefore crucial for the long-term survival and adaptive potential of wild populations. Together with the growing recognition of genetic diversity as the foundation for ecosystem protection and ecological restoration (*e.g.,* (Ngeve, 2025)), our results highlight the urgent need to integrate genetic impacts into chemical risk assessment and, conversely, to account for chemical exposure in conservation planning. Because genetic changes can precede more severe ecological effects of chemical pollution (*e.g.,* species loss), further development and validation of genetic based methods could complement traditional species-level impact measures (*e.g.,* species sensitivity distributions). With more comprehensive genetic and chemical data, the landscape genetics approaches could also help identify populations, species, or regions at heightened risk and, following independent validation, contribute to early-warning approaches for detecting biological responses to chemical exposure in wildlife. More broadly, the workflow developed in this study could be applied to other non-target wildlife species for which comparable datasets exist. Such data-reuse approaches could complement conventional environmental monitoring and contribute to broader One Health frameworks linking chemical exposure, wildlife health, and ecosystem health and represent a starting point for incorporating genetic impacts of chemical exposure into environmental risk assessment.

## Data Availability Statement

Input data were obtained from refs (Iacolina *et al*., 2016, 2018; de Jong *et al*., 2023; Pistocchi *et al*., 2023). Results tables are added as supplementary material.

## Acknowledgements

This work has received funding from the Swiss State Secretariat for Education, Research and Innovation (SERI). This research is part of the Horizon Europe TerraChem project and was funded by the European Union (Project number 101135483).

## Declaration of generative AI and AI-assisted technologies in the manuscript preparation process

During the preparation of this work, the author(s) used AI-based assistance (OpenAI, GPT) to support text structuring, clarity, and language refinement, as well as code restructuring. After using this tool, the authors reviewed and edited the content as needed and take full responsibility for the content of the published article.

