## Supplemental figures for "Integration of publicly available datasets reveals pesticide-associated genetic variation in a wild non-target animal"

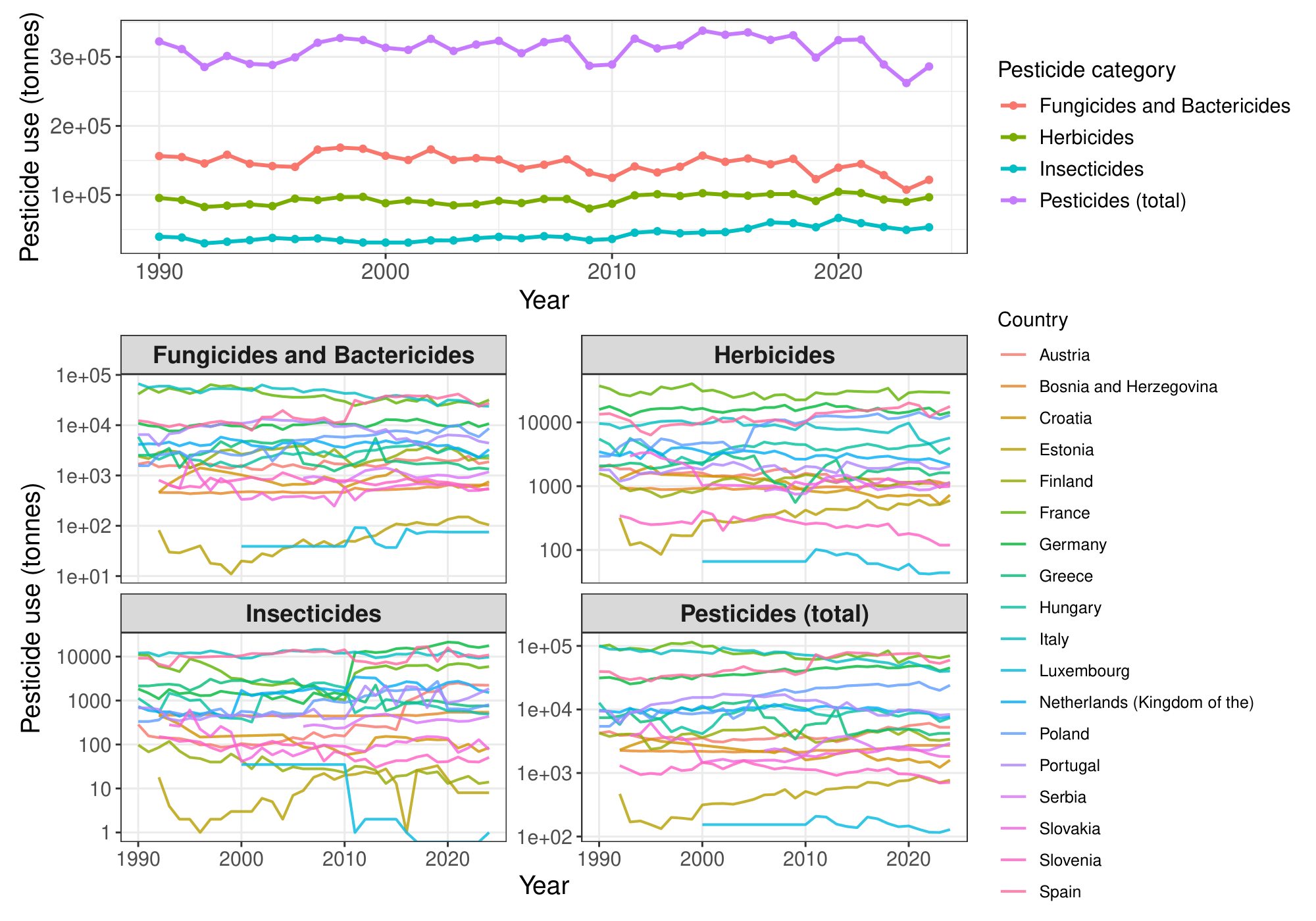

**Figure S1. Temporal trends in pesticide use across European countries.**

(A) Annual pesticide use summed across all included countries for each pesticide category. (B) Country-specific trends in pesticide use, with each line representing an individual country. Note that the y-axis in panel B is displayed on a log10 scale. Pesticide use is expressed in tonnes. Data were obtained from FAOSTAT.

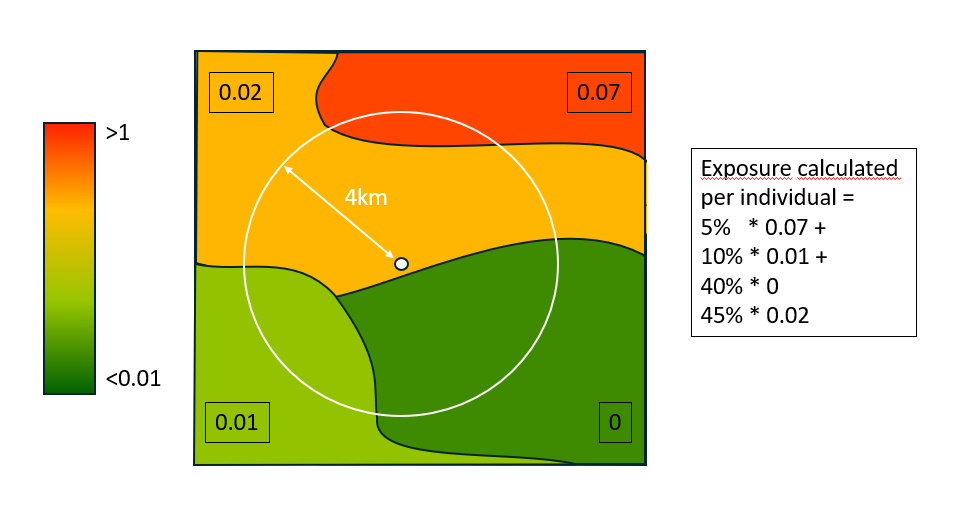

Exposure calculated per individual =

5% * 0.07 +

10% * 0.01 +

40% * 0 +

45% * 0.02

**Figure S2. Pesticide concentration calculation per individual.**

This figure is intended solely to demonstrate the calculation approach. It does not contain or represent real data.

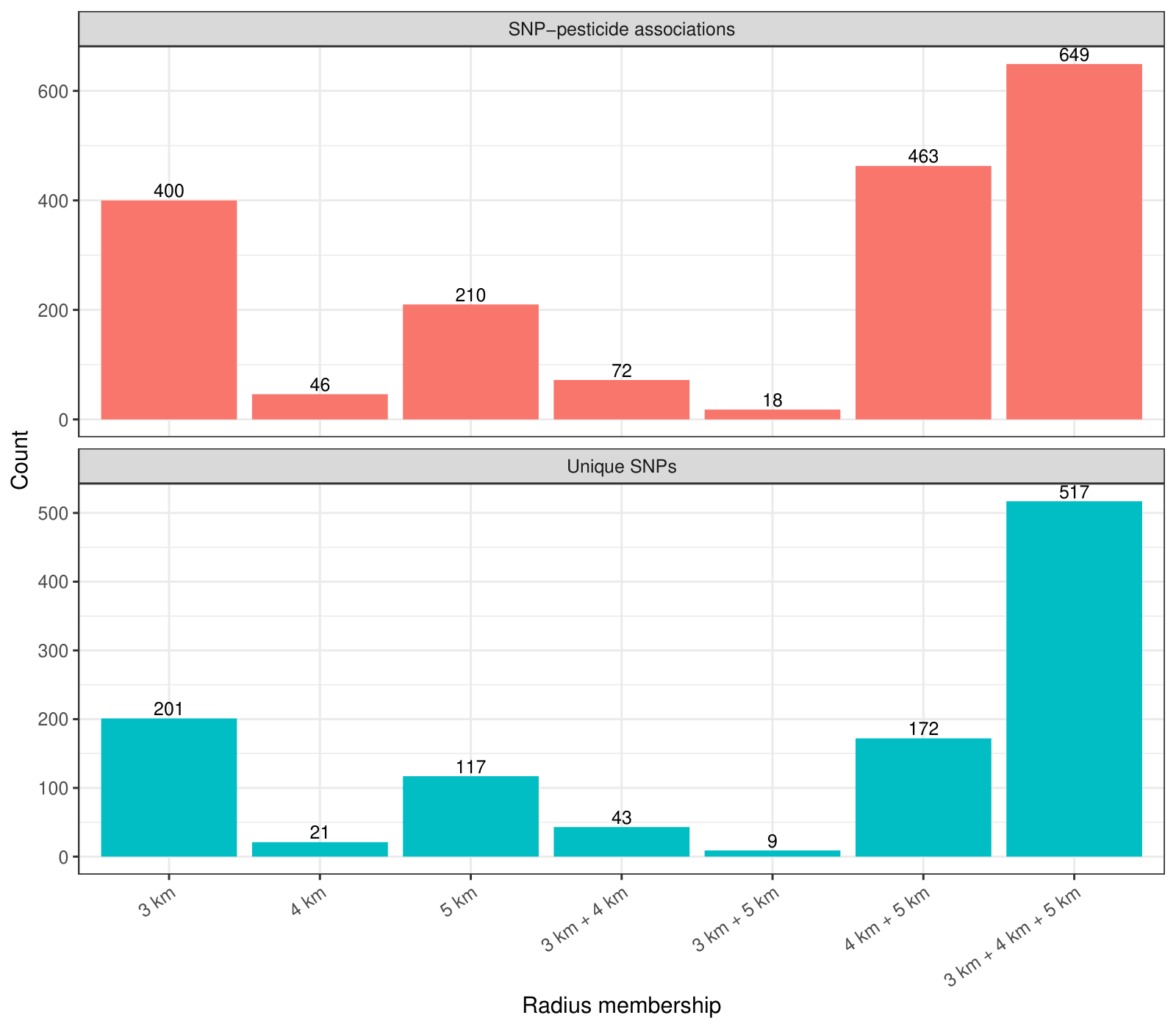

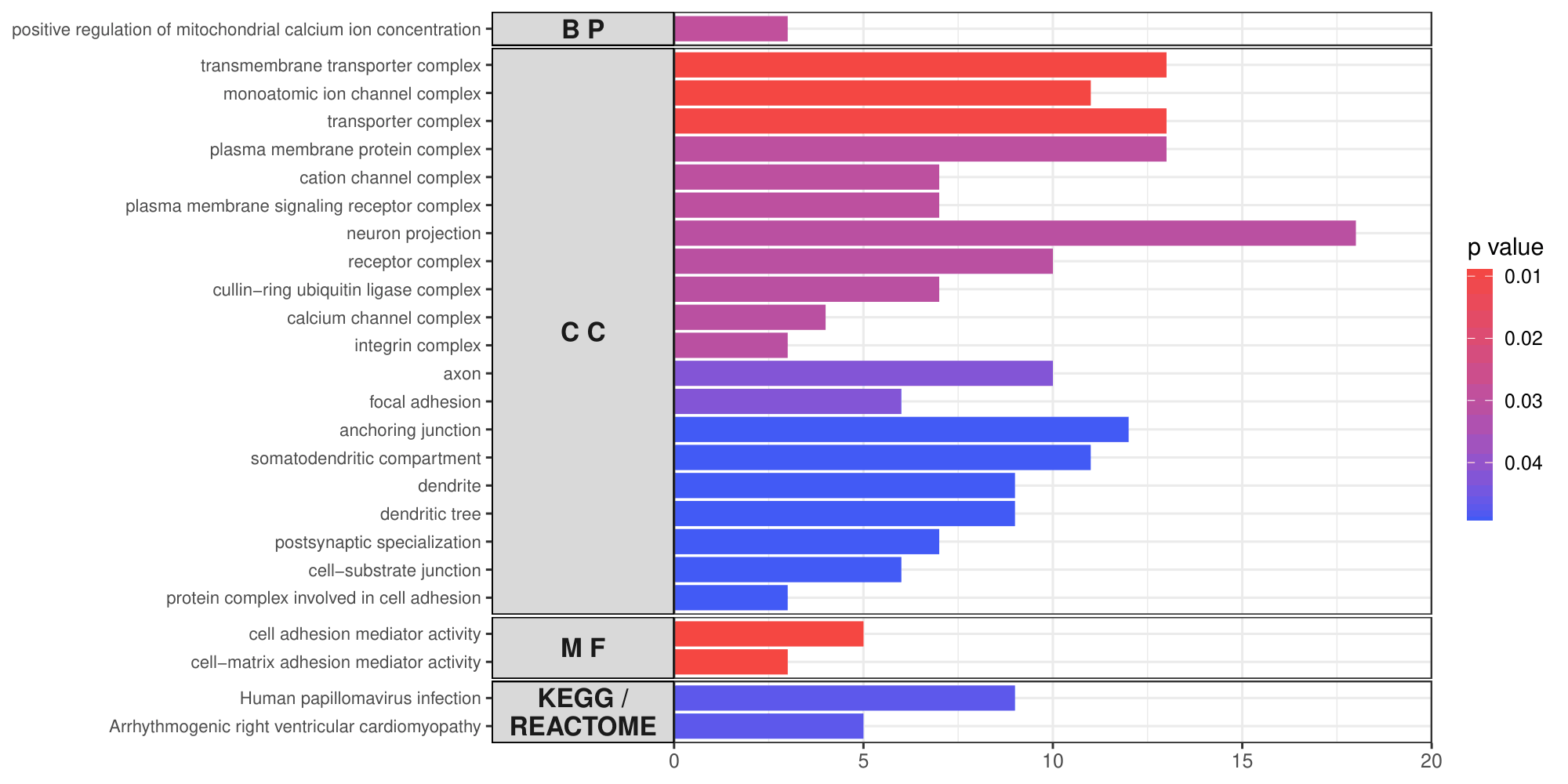

**Figure S3. Overlap across three spatial scales**

**Pesticide-associated SNPs (top panel).** Number of unique and shared SNPs and SNP–pesticide associations identified using pesticide concentrations averaged within 3-, 4-, and 5-km radii around each wild boar sampling location. Bars show associations unique to each radius or shared between two or all three radii. **Functional enrichment (bottom panel).** Enriched Gene Ontology (GO) terms and KEGG/Reactome pathways for genes consistently identified across the three radii. Bars indicate the number of overlapping genes associated with each enriched term, with colour representing the enrichment p-value. BP, biological process; CC, cellular component; MF, molecular function.

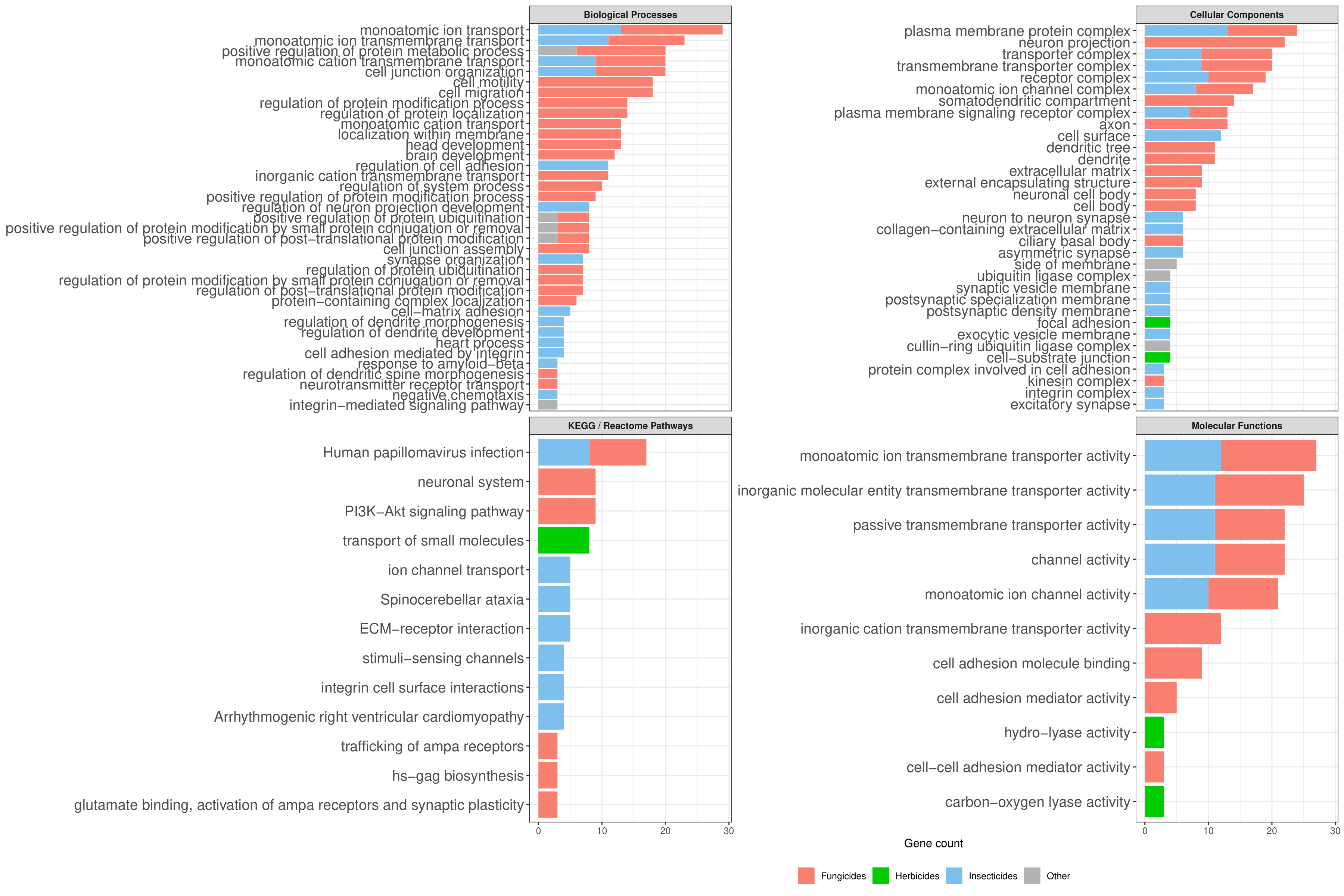

**Figure S4. Enrichment results for pesticide classes**

Enriched Gene Ontology (GO) terms and KEGG/Reactome pathways identified after pooling genes associated with pesticides within each pesticide class (insecticides, herbicides, fungicides, and other pesticides). Bars indicate the number of genes overlapping with each enriched term and colours indicate pesticide class.

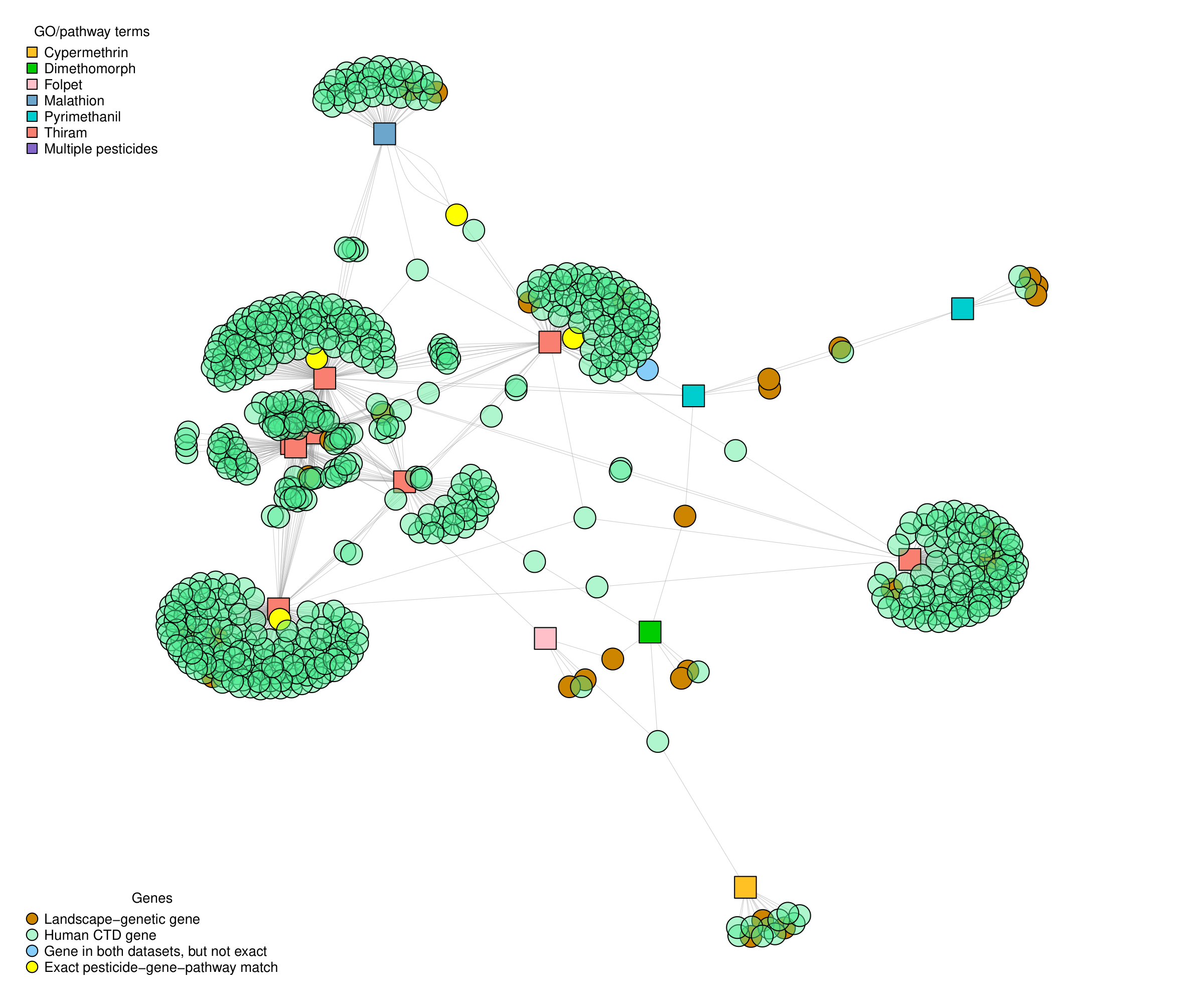

**Figure S5. Network diagrams showing external support for human Gene Ontology and pathway terms with SNPs associated with environmental pesticide concentrations**

Network diagram of GO terms enriched for pesticides in our landscape genetics study with external support in GO/pathway terms enriched for CTD *Homo sapiens* chemical – gene associations. Different genes were significant between our analysis and CTD (shown by green and orange circles) and some genes overlapped (yellow circles). Squares are GO terms associated with each chemical (color-coded by chemical) and circles are genes enriched by each chemical in each GO term (color-coded by the source of the gene). A line connecting a GO term and a gene means that gene was enriched in that GO term.
